# Drone phenotyping and machine learning enable discovery of loci regulating daily floral opening in lettuce

**DOI:** 10.1101/2020.07.16.206953

**Authors:** Rongkui Han, Andy J.Y. Wong, Zhehan Tang, Maria J. Truco, Dean O. Lavelle, Alexander Kozik, Yufang Jin, Richard W. Michelmore

**Affiliations:** The Genome and Biomedical Sciences Facility, University of California, Davis; Department of Land, Air and Water Resources, University of California, Davis

**Author notes:** **Author Contributions:** RM, MJT, DL, and RH conceived the experiment. RH designed and conducted the field experiment, performed the data analysis including the machine learning, Bayesian inference, and genetic mapping, and drafted the paper. AW, ZT, and YJ performed the drone phenotyping and assisted in the image analysis. MJT and DL developed the mapping population and genotyped by sequencing. AK made the video of asynchronous flower opening. All authors contributed to writing the paper.

**Keywords:** flowering, flower opening, genetic mapping, QTL mapping, lettuce, drone, unmanned aerial vehicle (UAV), high-throughput phenotyping, remote sensing phenotyping, image analysis, machine learning, support vector machine (SVM), Bayesian inference

## Abstract

Flower opening and closure are traits of reproductive importance in all angiosperms because they determine the success of self- and cross-pollination. The temporal nature of this phenotype rendered it a difficult target for genetic studies. Cultivated and wild lettuce, *Lactuca spp*., have composite inflorescences comprised of multiple florets that open only once. Different accessions were observed to flower at different times of day. An F_6_ recombinant inbred line population (RIL) had been derived from accessions of *L. serriola* x *L. sativa* that originated from different environments and differed markedly for daily floral opening time. This population was used to map the genetic determinants of this trait; the floral opening time of 236 RILs was scored over a seven-hour period using time-course image series obtained by drone-based remote phenotyping on two occasions, one week apart. Floral pixels were identified from the images using a support vector machine (SVM) machine learning algorithm with an accuracy above 99%. A Bayesian inference method was developed to extract the peak floral opening time for individual genotypes from the time-stamped image data. Two independent QTLs, *qDFO2.1* (*Daily Floral Opening* 2.1) and *qDFO*8.1, were discovered. Together, they explained more than 30% of the phenotypic variation in floral opening time. Candidate genes with non-synonymous polymorphisms in coding sequences were identified within the QTLs. This study demonstrates the power of combining remote imaging, machine learning, Bayesian statistics, and genome-wide marker data for studying the genetics of recalcitrant phenotypes such as floral opening time.

**One sentence summary:** Machine learning and Bayesian analyses of drone-mediated remote phenotyping data revealed two genetic loci regulating differential daily flowering time in lettuce (*Lactuca spp.*).

## Introduction

Floral opening is a complex and dynamic process marked by rapid, drastic changes in the morphology of the reproductive organs of angiosperms. The time of floral opening marks the onset of the period during which cross pollination becomes possible, making this physiological process a critical phase in plant sexual reproduction. From an ecological perspective, different floral opening times within the day can play an important role in population divergence by contributing to temporal reproductive isolation (Matsumoto et al., 2013). Synchronizing floral opening time with peak activity of effective pollinators may help improve outcrossing and reproductive success (Sakamoto et al., 2012).

Across different flowering species, successful floral opening is accomplished through a diverse set of events—petals may unfold, spiral outward, or spring open, depending on their particular anatomies, and the opening process may or may not be reversible. What usually underlies these impressive local movements is a high rate of cell expansion and/or abscission driven by changes in osmotic pressures. The timing of this process is regulated by external and internal factors. Environmental cues, such as humidity, temperature, and light, the internal circadian rhythm of the plant, and hormone signaling all modulate floral opening (van Doorn and van Meeteren, 2003; van Doorn and Kamdee, 2014). Different species show various levels of responsiveness to these internal and external cues. In extreme cases, the effect of the same signal can be completely opposite in different species. For instance, ethylene treatment is known to accelerate floral opening in some rose (*Rosa spp.*) cultivars, while inhibiting floral opening in others (Reid *et al.* 1989).

The molecular control of floral opening is incompletely understood. Past endeavors to probe regulation of floral opening have mainly taken four approaches: gene transcription, cellular signaling, mutant analysis, and forward genetics. Transcription-level events corresponding to the physiological process of floral opening have been detected in multiple studies. High accumulation of volatile-emission-related R2R3-MYB transcription factor EOBII was found in hybrid peas (*Pisum x hybrida* “Mitchell Diploid”) and *Nicotiana attenuata* prior to floral opening. RNAi knockdown of EOBII resulted in failure to enter anthesis and premature senescence (Colquhoun et al., 2011). Over-expression of fructan 1-exohydrolase was associated with flower opening in *Campanula rapunculoides,* presumably contributing to decreasing osmotic pressure in expanding petals by breaking down polysaccharide fructan (Vergauwen et al., 2000; Le Roy et al., 2007). Similarly, transcriptional upregulation of cell-wall-loosening expansin was associated with floral opening in carnation (*Dianthus caryophyllus*; Harada et al., 2010). Transcription-level fluctuation of ethylene receptors during flowering was reported in tree peony (*Paeonia suffruticosa*; Zhou et al., 2010). Phytochrome activity is also involved. Kaihara and Takimoto (1980) demonstrated that a flash of red light during the night before anticipated floral opening can alter the time of floral opening on the following day. The effect of red light was diminished by a subsequent exposure to far-red light. MicroRNA regulation of flower opening was proposed after comparing microRNA levels between buds and flowers in 5-year-old plum blossom trees (*Prunus mume*; Wang et al., 2014). Nevertheless, the regulatory network that oversees transcription alteration remains unclear (van Doorn and Kamdee, 2014). Few mutants specific to floral opening have been identified in model plant systems (van Doorn and Kamdee, 2014); a mutation in a RINGv E3 ubiquitin ligase that causes reduced cutin biosynthesis or loading was found to cause a lack-of-opening phenotype in oilseed rape (*Brassica napus*) suggesting the critical role of cutin in successful floral opening (Lu et al., 2012). Only one forward genetic study has investigated the genetic regulation of floral opening time (Nitta et al., 2010); the segregation of morning flowering versus evening flowering in an F_2_ population derived from a hybrid between daylily (*Hemerocallis fulva*) and night lily (*H. citrina*) suggested the presence of a major effect gene. This study also suggested independent regulation of floral opening and closure times in lily.

In order to understand more about the genetic regulation of floral opening time, we analyzed natural variation in this phenotype in *Lactuca serriola* (wild lettuce) and *L. sativa* (lettuce). *Lactuca* spp. are members of the Compositae family with compound hermaphrodite inflorescences that only open once. *L. serriola* is the wild progenitor of modern cultivated lettuce and is fully reproductively compatible with *L. sativa.* We took advantage of a recombinant inbred line (RIL) population developed from a cross between accessions of *L. serriola* and *L. sativa* that differed for floral opening time by 3.5 hours. We overcame the challenge of studying floral opening time in a large, replicated RIL population by utilizing drones equipped with a multi-spectral camera to repeatedly image the entire experimental field. Effectiveness of drones in high-throughput crop phenotyping has been demonstrated in recent studies (Spindel et al., 2018; Xu et al., 2019). In our study, data from hourly drone flights were analyzed using an innovative combination of machine learning and Bayesian statistics to quantify the floral opening phenotype. Two significant quantitative trait loci (QTLs) collectively explained more than 30% of the phenotypic variation of floral opening time; these QTLs contained genes known to regulate circadian rhythms in Arabidopsis.

## Results

Most lettuces, including the oil seed type PI251246, start to flower early in the morning. In contrast, *L. serriola* accession Armenian999 does not flower until the afternoon. Two-hundred and thirty-six F6 RILs that had been developed from crossing these two accessions were available for investigating the genetic basis of asynchronous floral opening phenotype. This phenotype is illustrated in a short video made using time lapse photography of two RILs from this population (https://www.youtube.com/watch?v=9w8iRTHXBxM) taken from an experimental field in Davis, CA, in June 2014, which corresponds to a 3-hour span in real time. In this video, flowers of one RIL begin to open approximately 55 minutes before the other. This segregating phenotype is also illustrated by photographs of individual flowers taken over an 8-hour time span of four RILs and the parents grown in a screenhouse in Davis, CA in June 2020 (Figure 1).

**Figure 1.**
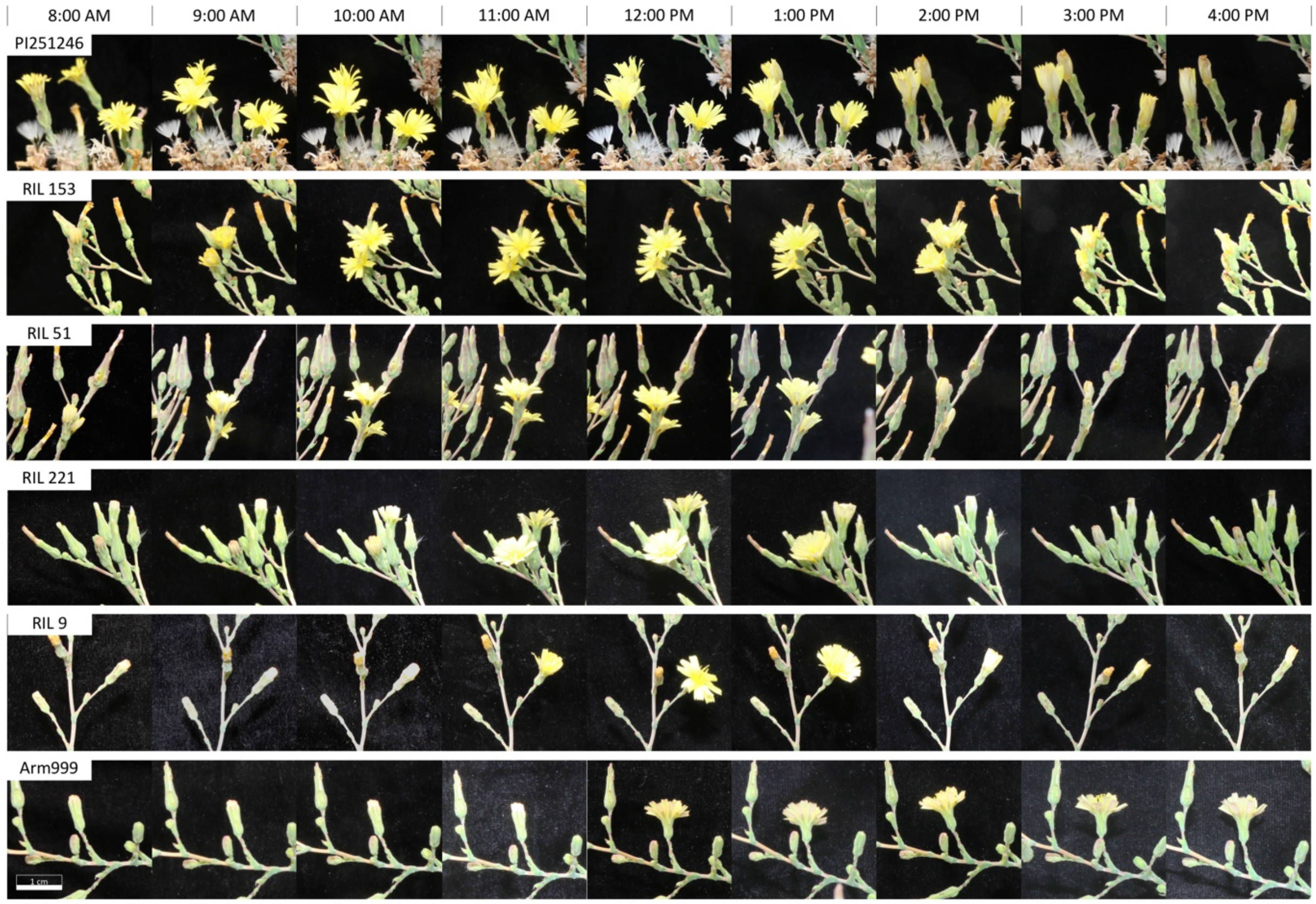
Close-up photographs taken at hourly intervals from 8.00 am to 4.00 pm illustrating the asynchronous floral opening and closing of the parental lines and four RILs of the PI251246 x Armenian999 F_6_ population.

### Remote sensing phenotyping

The 236 RILs, both parental lines, and two controls, *L. sativa* cv. Salinas and *L. serriola* accession US96UC23, were planted in two complete randomized replicates of eight plants in Davis, CA, during summer 2019. Multi-spectral images were captured at 9 am, 11 am, 1 pm, and 3 pm on July 1^st^, 2019, and 10 am, 12 pm, 2 pm, and 4 pm on July 9^th^ using a multispectral camera mounted on a drone. Each drone flight took an average of nine minutes. On average, 2,309 raw images with 85% front- and side-overlap were generated per flight. The 2 pm drone flight did not generate any images due to technical errors and had to be discarded. A total of seven whole-field images at 1 cm spatial resolution were generated by GPS-guided tiling of raw images. Yellow floral pixels are visible to the naked eye from whole-field images (Figure 2 and Supplementary Figure S1).

**Figure 2.**
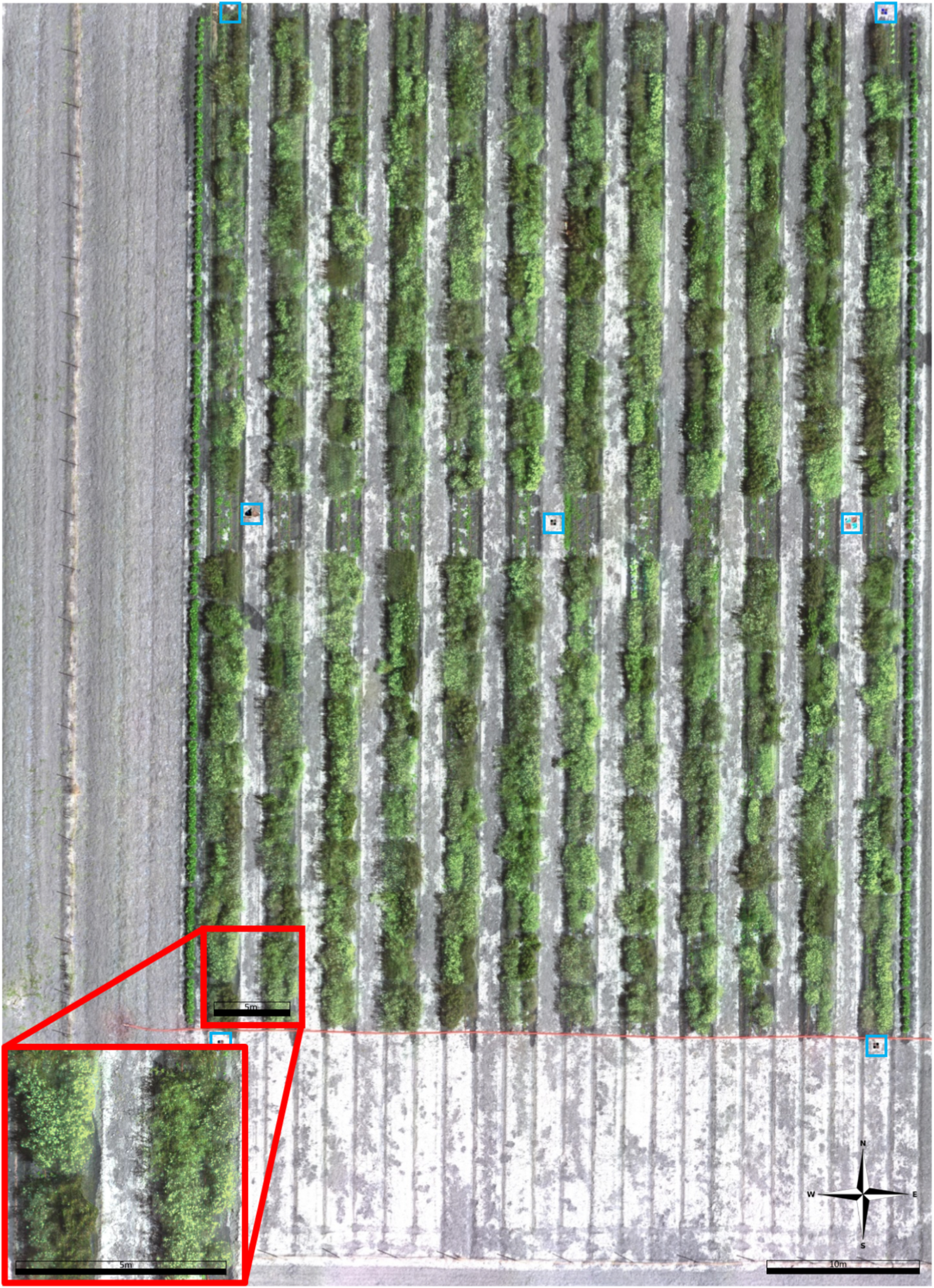
Reconstructed field orthomosaic image at 11:00 am on July 1^st^, 2019. A total of 2,355 raw drone images in five reflectance bands (red, blue, green, red-edge and infrared) were tiled using the Pix4Dmapper Pro software in GPS-guided mode. Only the red, blue and green channels are shown in this figure. The seven ground control points are identified with blue boxes.

### Machine learning classification of floral pixels

A total of 4,807 floral, vegetative, and ground pixels were randomly sampled from the super-high-resolution field images at all seven time points (Supplementary Table S1). Pairwise scatterplots of the Hue-Saturation-Value (HSV) values of the sample pixels indicated clear distinction between pixels belonging to different categories (Figure 3a). Images taken at different time points appeared to be largely homogeneous regarding pixel HSV (Figure 3b).

**Figure 3.**
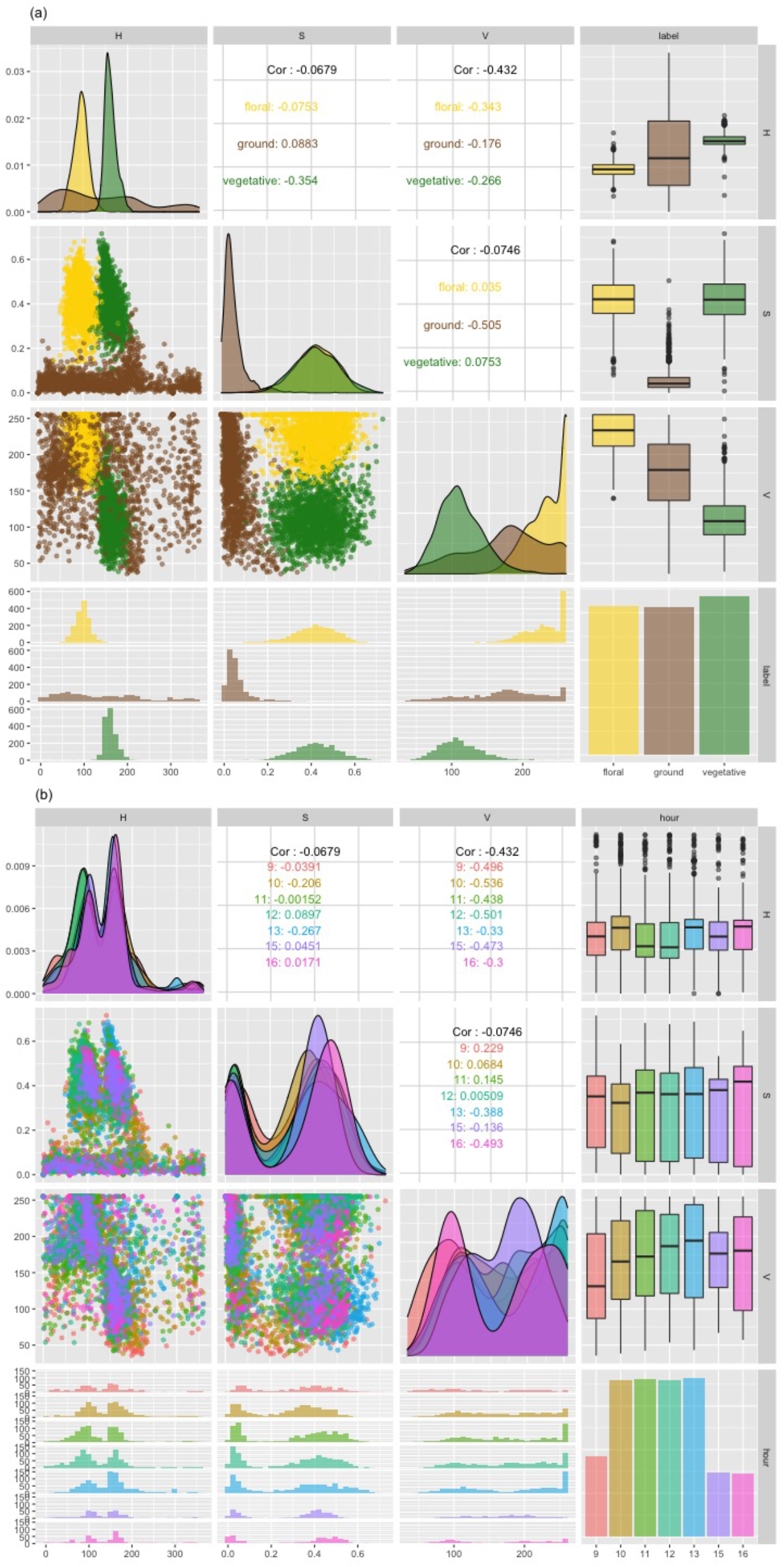
Pair-wise distribution of hue (H), saturation (S) and value (V) of sampled floral (yellow), ground (tan), and vegetative (green) pixels. The plots are colored by pixel labels in (a) and hours in (b). Fewer samples were taken at 9 am, 3 pm and 4 pm due to the small number of floral pixels available at these times. 2(a) demonstrates clear distinction between the three classes of pixels; 2(b) indicates that images taken at different hours of the day are homogeneous in their composition.

Machine learning was used to classify pixels from whole-field images into “floral,” “vegetative,” or “ground” categories. The 4,807 labeled sample pixels were divided evenly into a training dataset (2,404 samples) and a testing dataset (2,403 samples). Five machine learning algorithms were trained using the training dataset. Ten-fold cross-validation test of pixel classification accuracy showed that the support vector machine (SVM) model outperformed all other models with a mean classification accuracy of 99.15%. When used to classify pixels in the testing dataset, the SVM model made predictions with 99.08% accuracy (Table 1 and Figure 4b). A final SVM model was trained using all sample pixels and tuning hyperparameters sigma = 1.102 and C = 0.5. The final model produced a total of 308 support vectors and had within-sample prediction accuracy of 99.12%.

**Figure 4.**
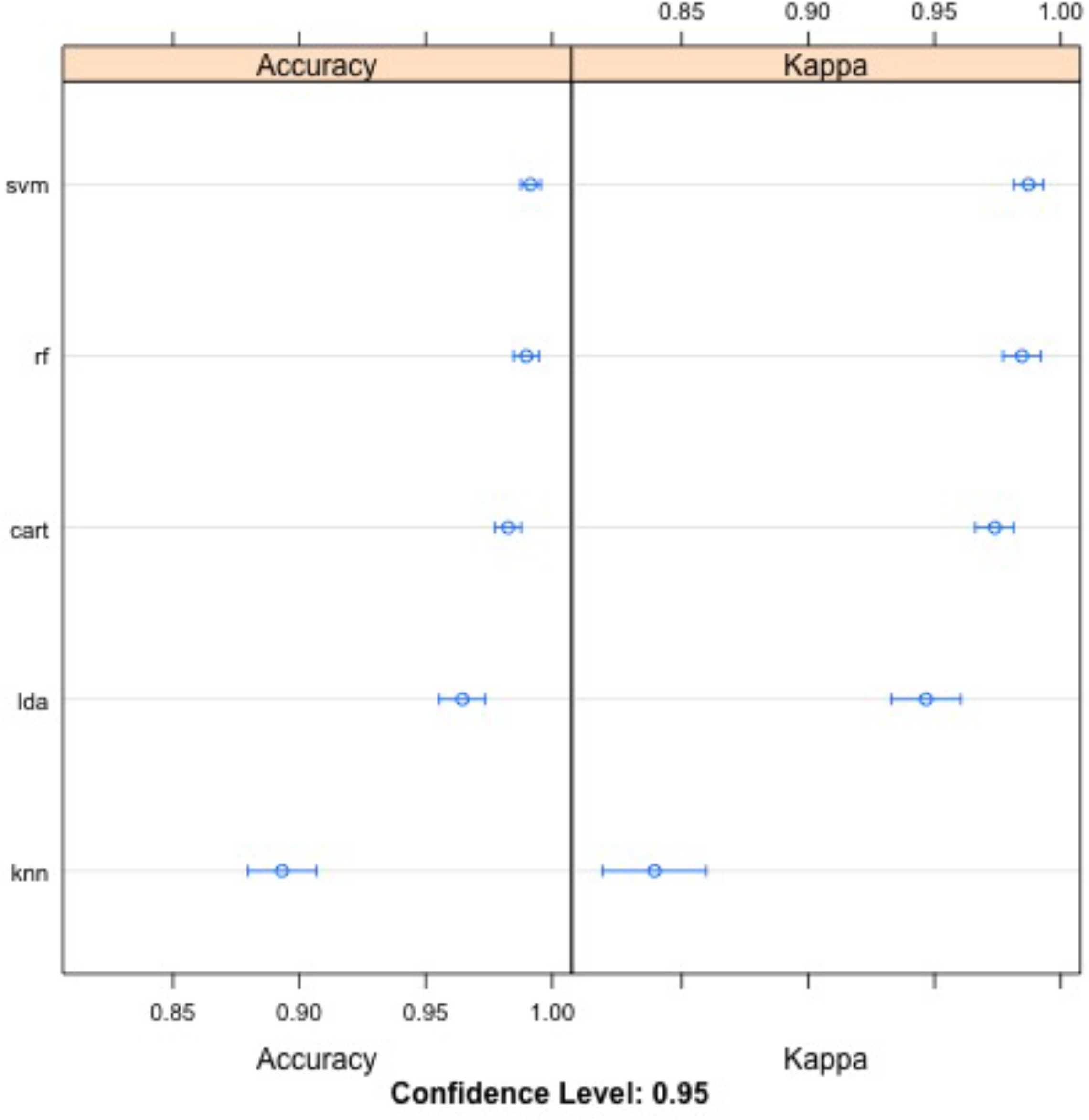
Comparison of pixel classification prediction accuracy and adjusted accuracy (“Kappa”) of five machine learning algorithms: support vector machine (SVM), random forest (RF), classification and regression tree (CART), linear discriminant analysis (LDA) and k-nearest neighbors (KNN).

**Table 1.** Prediction accuracy of support vector machine algorithm, checked using testing dataset

|  | Testing: floral | Testing: vegetative | Testing: ground |
| --- | --- | --- | --- |
| Prediction: floral | 792 | 1 | 4 |
| Prediction: vegetative | 3 | 779 | 8 |
| Prediction: ground | 7 | 6 | 803 |

The final SVM model was deployed to predict floral pixels in all field images. Plot-level floral pixel counts at all seven time points are listed in Supplementary Table S2. The total number of predicted floral pixels ranged from 2,730 at 4 pm to a daily maximum of 1,395,676 at 11 am. The change in the total number of floral pixels throughout the course of the day peaked in the late morning, consistent with maximum floral opening on a population level being between 11 am and 12 pm (Figure 5).

**Figure 5.**
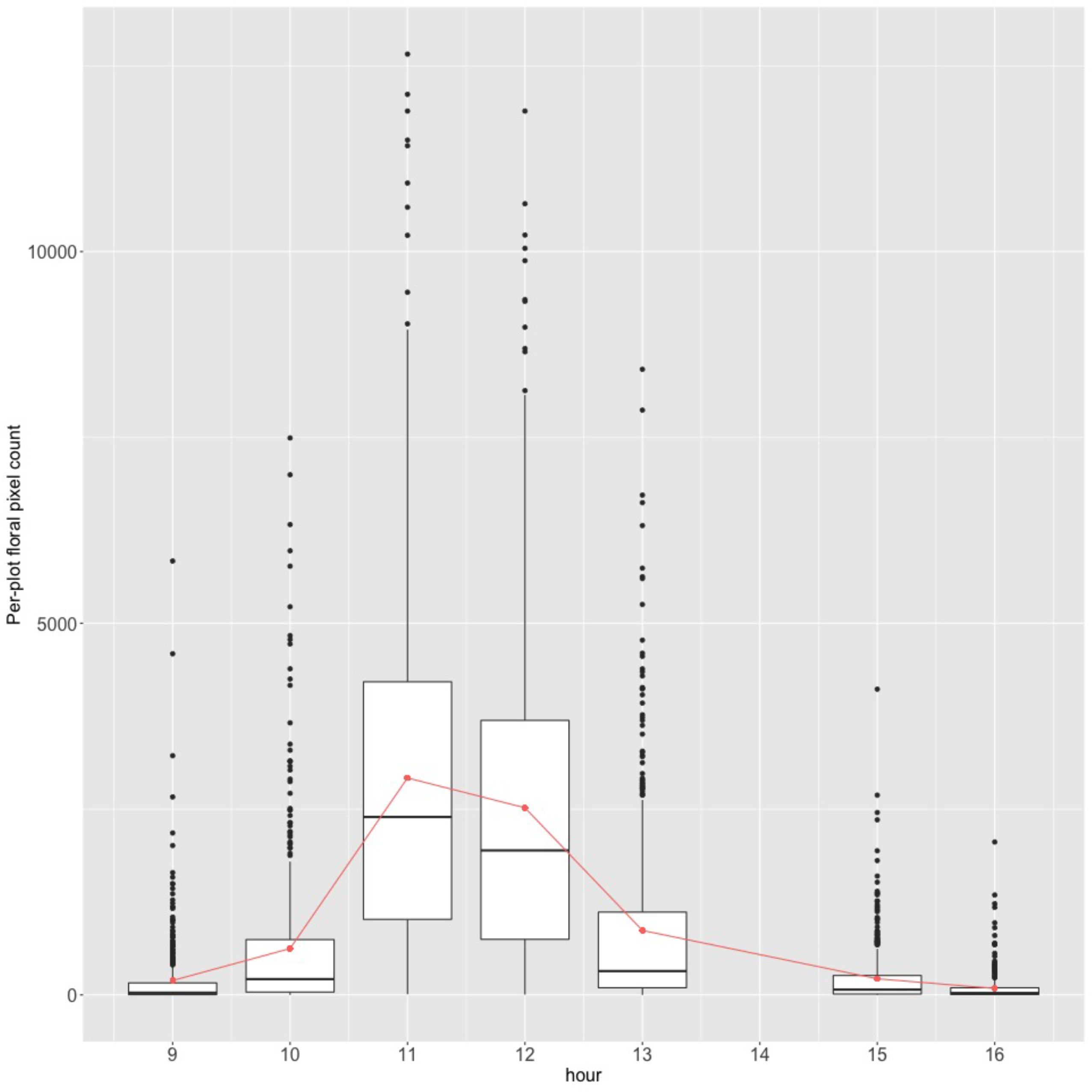
The ranges of per-plot number of floral pixels at each time point. The orange line shows the change of the mean per-plot floral pixel count over time. The box plots show median and quartile of the pixel count distribution at each time point.

### Bayesian inference of peak floral opening time

The peak floral opening time (FOT) of individual plots was inferred from the hourly floral pixel count for each plot. Figure 6 is a visualization of the output of SVM floral pixel classification. Plot “1” in the blue box clearly reached peak opening early in the day, near 10 am. Similarly, plot “2” in the green box had peak FOT near 11 am, plot “3” (yellow box) near 12 pm, and plot “4” (orange box) near 1 pm (Figure 6). The temporal increase and decrease of the floral pixel count throughout the course of a day within each plot can be described using a distinct bell-shaped curve, with parameter τ characterizing the peak FOT of the plot, i.e., the mean and mode of the curve, and parameter δ^2^ characterizing the duration of the opening within the plot.

**Figure 6.**
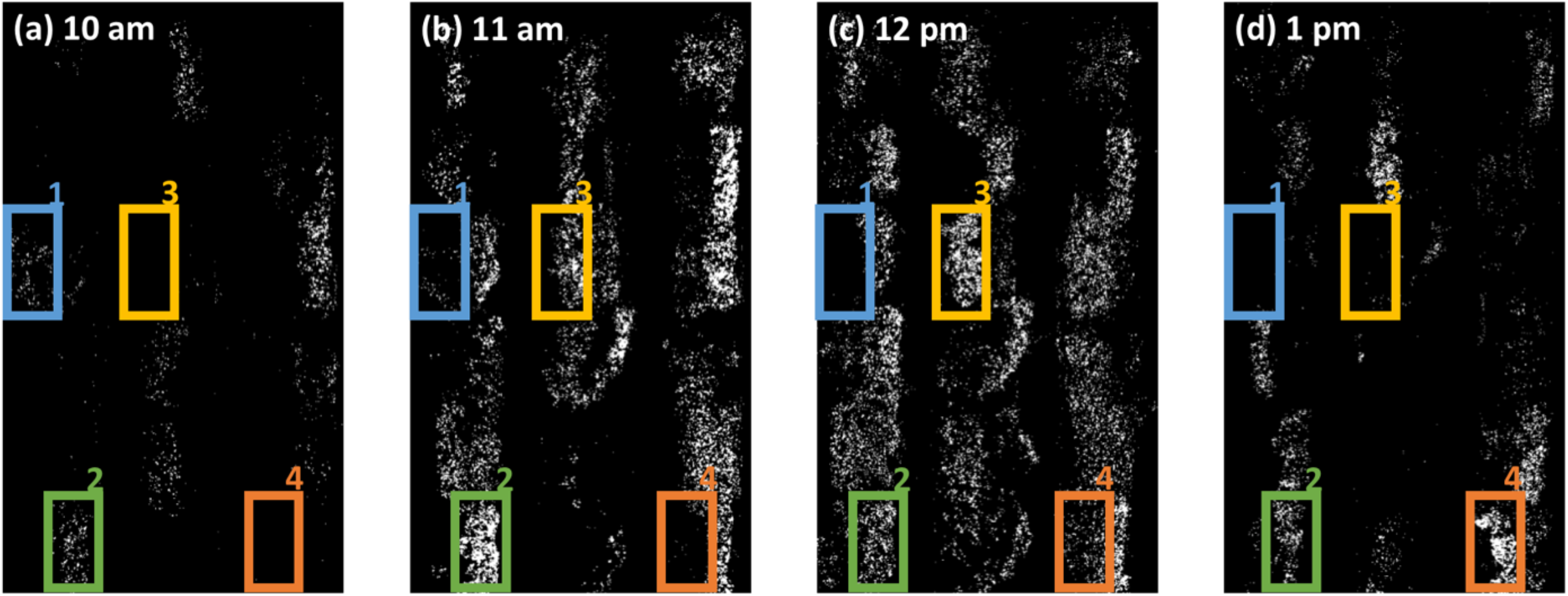
False-colored image of 36 plots based on output of SVM classification of image pixels. Floral pixels were rendered white and nonfloral pixels (vegetative or ground) were illustrated in black. Changes in floral pixel count of highlighted plots “1”, “2”, “3”, and “4” through hours (a) 10 am, (b) 11 am, (c) 12 pm and (d) 1 pm show the variation in floral opening time.

A bell-shaped likelihood function with parameters {τ_k_, δ_k_^2^} that best described the hourly floral pixel fluctuation of plot k (k = 1, 2, …, 480) was fitted to the time-series floral pixel data for each plot using Markov Chain Monte Carlo (MCMC). Divergent incidences were re-fitted by the MCMC to account for possible poor initialization. All plots converged after two iterations of sampling. The inferred peak FOT between the two blocks showed strong correlation (R^2^ = 0.485; Figure 7, Figure 8, and Supplementary Figure S2). Simple linear regression model reported no significant difference of peak FOT between the two blocks (p = 0.10). Therefore, the mean phenotype was calculated using the simple Euclidean mean between the two blocks.

**Figure 7.**
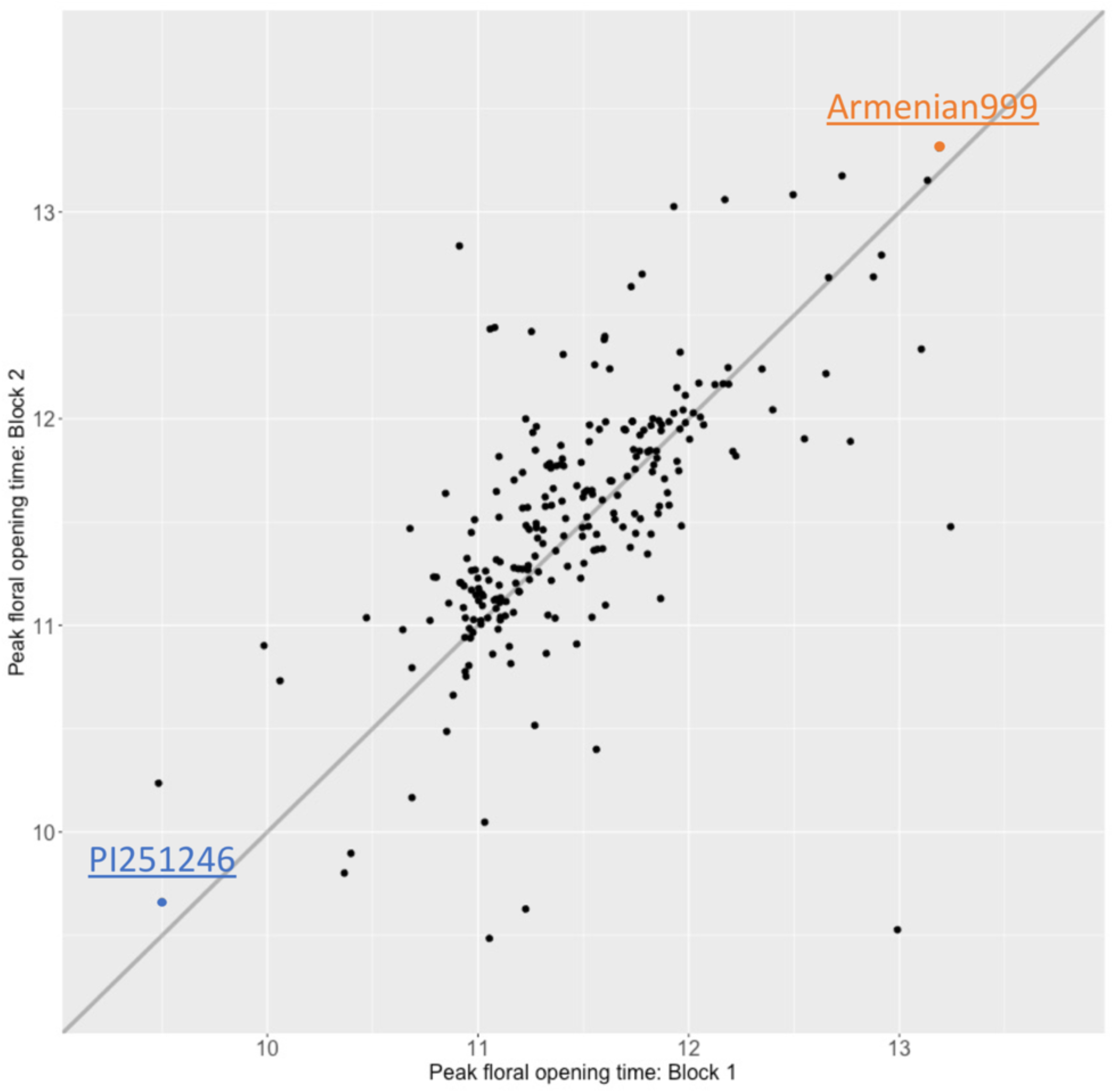
Correlation between-blocks for peak floral opening time of 236 RILs that had a convergent inference and an estimate for both blocks (R^2^ = 0.485).

**Figure 8.**
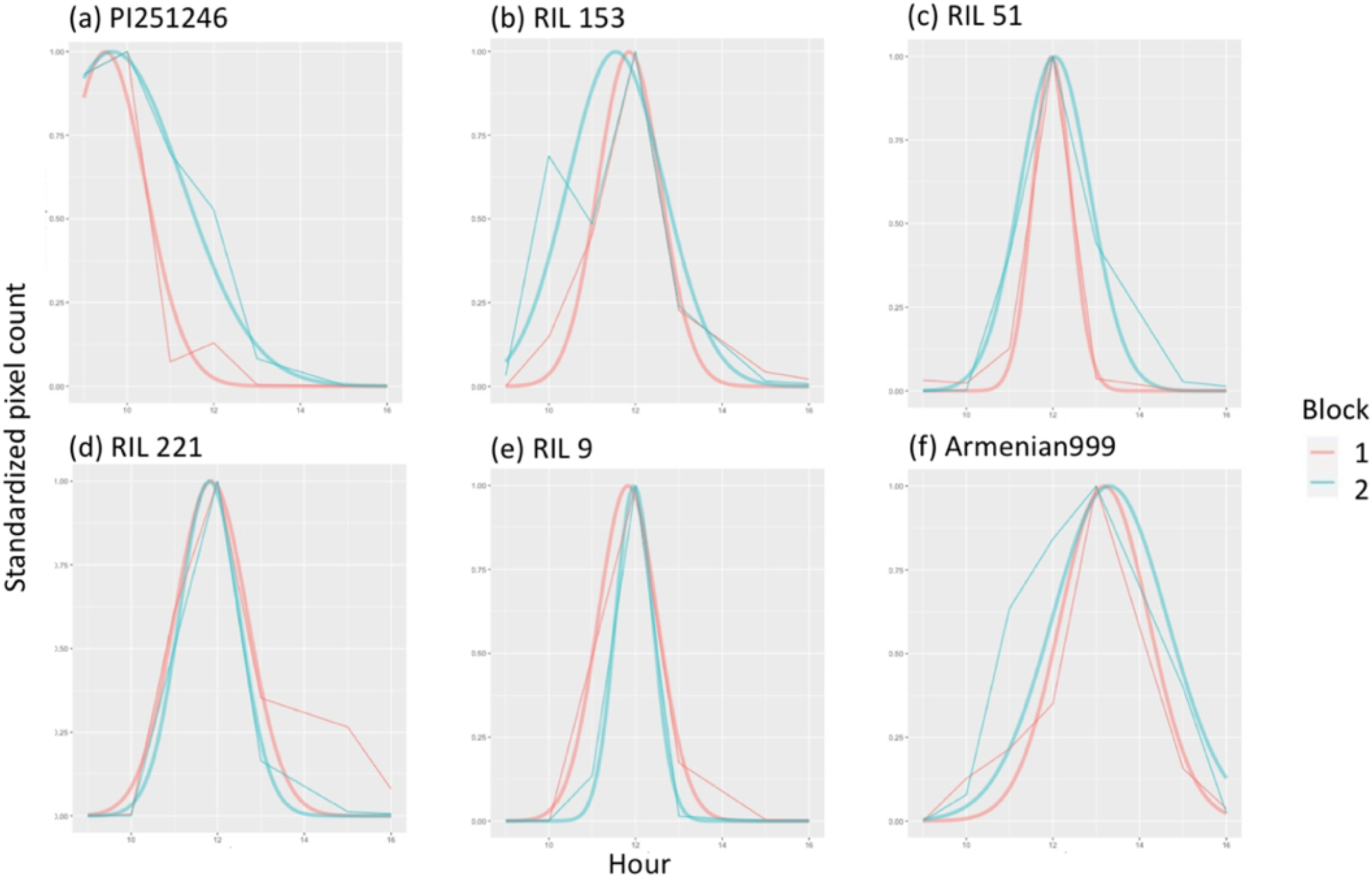
Standardized floral pixel counts for four RILs and the parents throughout the day, overlaid with the respective Bayesian inferred floral opening curve. Close-up photographs of floral opening and closing events of these RILs and the parents are shown in Figure 1.

Inferred peak FOT ranged from 09:17 am to 1:08 pm, following a slightly heavy-tailed normal distribution with a mean of 11:29 am (Figure 9; Supplementary table S3). The standard deviation of phenotypic distribution is 36.8 minutes. The inferred peak FOT of the early opening parent, PI251246, was 9:34 am, while that of the late opening parent, Armenian999, was 1:15 pm. No obvious transgressive segregation was found within the population.

**Figure 9.**
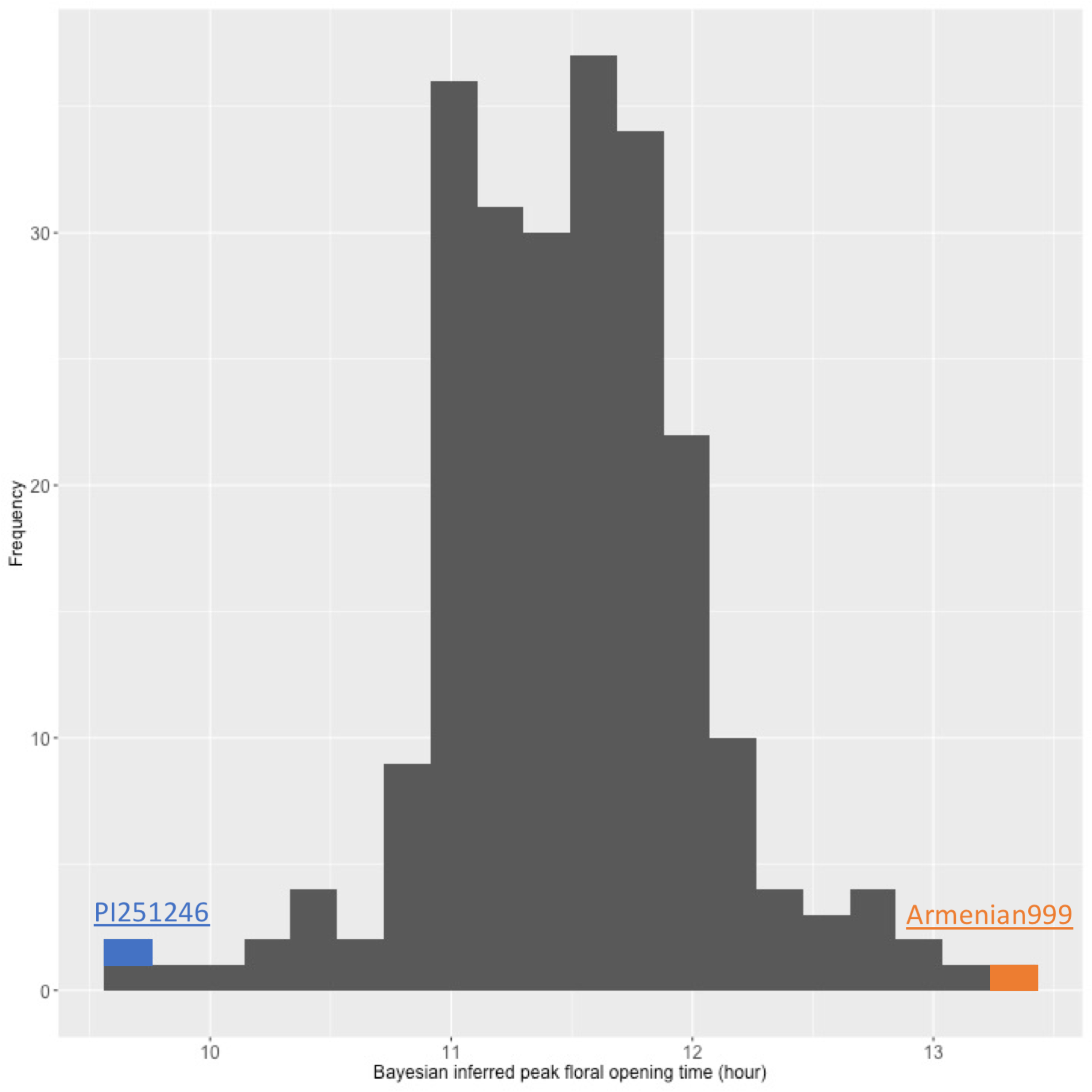
Distribution of high-confidence Bayesian inferred peak floral opening time of the parents and the 236 RILs used in QTL the analysis.

### Genotyping

A total of over 354 million 100 bp Illumina reads were obtained from all RILs. Mapping reads to version 8 of the lettuce reference genome (Reyes-Chin-Wo et al., 2017) yielded 422,418 single nucleotide polymorphism (SNP) markers, covering all nine chromosomes of the lettuce genome. After filtering against missing data and segregation distortion, 18,805 SNP markers remained. LepMap3 (Rastas, 2017) was used to produce a genetic map comprising 17,402 SNP markers in 2,677 genetic bins, covering 1,883 cM in the nine chromosomal linkage groups (Supplementary Figure S3), which is similar to the previously reported genetic map size (Truco et al., 2013). The heterozygosity rate of selected SNPs was 3.25%, consistent with the expected heterozygosity rate for F6 populations, 3.13%. No regions exhibited severe segregation distortion. One SNP marker was selected from each genetic bin, resulting in 2,677 markers for QTL mapping. The mean distance between each pair of adjacent markers is 0.7 cM. Four gaps larger than 5 cM are present in this map; these gaps are located at 149.0–155.8 cM on linkage group 3, 155.8–166.2 cM on linkage group 3, 62.8–68.1 cM on linkage group 7, and 50.3–55.7 cM on linkage group 9. Four RILs were excluded from downstream analyses due to the large percentage of missing genotype data, resulting in a final set of 232 RILs for QTL mapping.

### QTL analysis

Genotype and peak FOT phenotype data for 232 RILs were used for QTL mapping. Mixed effect modeling estimated the narrow sense heritability of the phenotype to be 0.8765. The significance threshold of the permutation test was set at negative log of odds (LOD) = 3.14 for the type I error rate of 0.05. Two significant QTLs were identified for peak FOT on Chromosomes 2 (LOD = 10.3) and 8 (LOD = 7.7) (Figure 10). The physical location of flanking markers and summary statistics of the effects of the two QTLs are detailed in Table 2. In both QTLs, the allele from the late-flowering parent, Armenian999, contributed to the later flowering phenotype (Figure 11).

**Figure 10.**
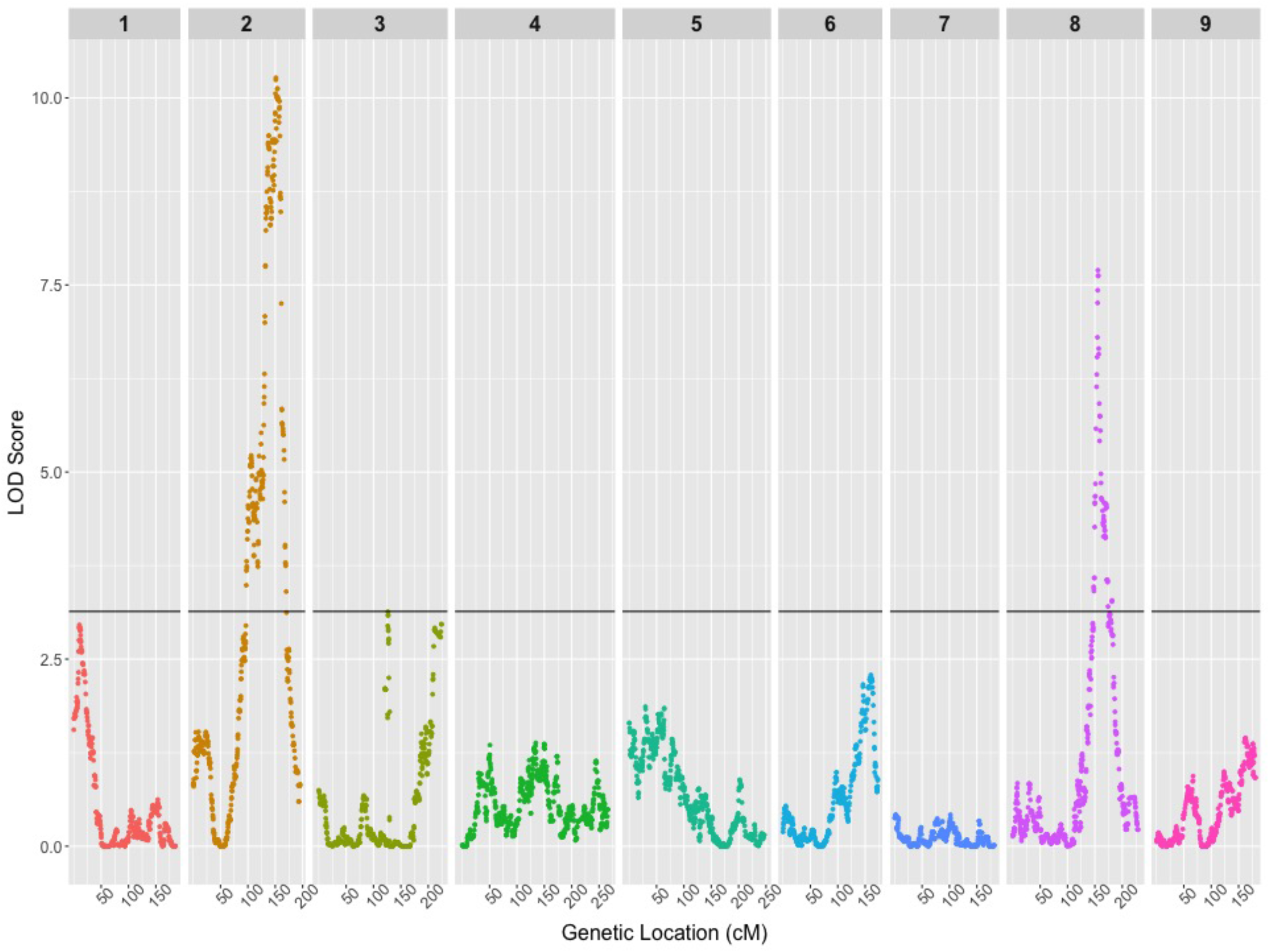
LOD scores of markers for peak floral opening time shown along the nine chromosomal linkage groups. The LOD threshold for significance (p < 0.05) calculated by 10,000 permutations is shown as a black line.

**Figure 11.**
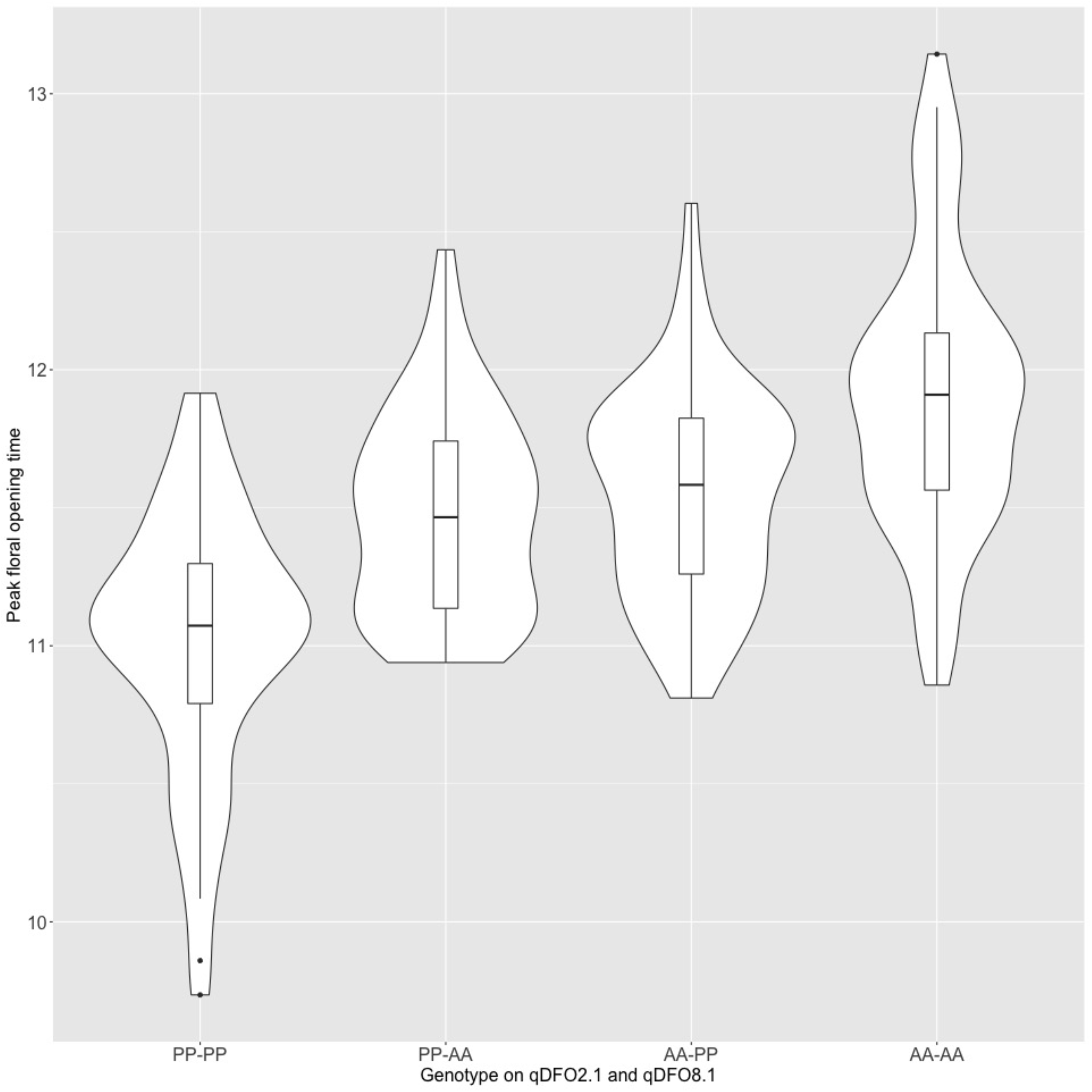
The additive effect of Armenian999 alleles at each QTL on peak floral opening time. The box plots represent the median and quartiles of the phenotypic distribution of each allelic combination. Widths of the violin plots represent density of samples at each phenotypic value.

**Table 2.** QTL linked to differential floral opening hour (*qDFO*) and their effects

| QTL | 1-LOD interval<br>Physical Location<br>(Base) | 1-LOD interval<br>Genetic<br>Location (cM) | LOD | p-value | Variance<br>Explained | Effect<br>of AA<br>allele<br>(hours) |
| --- | --- | --- | --- | --- | --- | --- |
| *qDFO2.1* | Chr2: 183,906,862–<br>190,964,979 | LG2: 148.7-159.0 | 10.3 | 0 | 18.23% | 0.48 |
| *qDFO8.1* | Chr8: 196,253,927–<br>202,987,597 | LG8: 153.4-156.9 | 7.7 | 0 | 13.84% | 0.4 |

### Candidate genes

The two significant QTLs, *qDFO2.1* (*Daily Floral Opening on chromosome 2)* and *qDFO8.1,* were investigated for candidate genes of known function in Arabidopsis. There are 309 gene models located within 1 LOD score on each side of the peak of *qDFO2.1* and 123 gene models within *qDFO8.1* in the reference annotation (Reyes-Chin-Wo et al., 2017). Among these 432 genes, 199 had coding sequence variants between the parents (Supplementary Table S4). Of the 1,752 orthologs of Arabidopsis genes involved in flowering time and/or circadian clock regulation, five were located within *qDFO2.1;* two of these genes exhibited coding sequence variants (Table 3). No orthologs involved in flowering time and/or circadian clock regulation were identified within *qDFO8.1*.

**Table 3.**
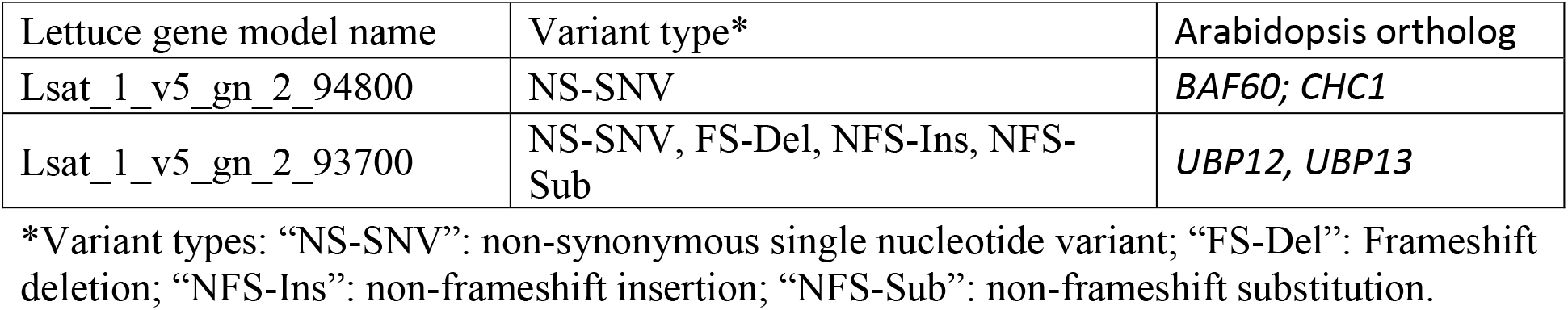
Candidate genes within QTL *qDFO2.1* with non-synonymous variations between parents and orthology to Arabidopsis genes involved in flowering time and/or circadian clock regulation.

## Discussion

Our study demonstrated the efficacy of using drones to detect quantitative temporal differences in floral opening events. We were able to collect multispectral data on a large number of plants multiple times per day. Machine learning enabled fast, robust recognition of phenotypes. Bayesian approaches provided a summary statistic for flower opening time of each line that was used for QTL analysis. This revealed two significant genomic regions determining floral opening time.

The machine learning algorithm predicted that the floral opening behavior of each single-genotype plot followed a Gaussian-like curve throughout the course of a day (Figure 5). The time-stamped floral pixel count data was then passed down to a Bayesian framework to extract summary statistics for floral opening time. The number of floral pixels captured by a drone image is a function of both the number of opening inflorescences and the degree of their opening (Figure 1). The peak floral opening time reflects the average opening time of all individual inflorescences within a plot. From an analytical standpoint, peak floral opening time is a better summary statistic for the floral opening process than the beginning or ending points because it is readily defined mathematically and is robust against detection errors. A Gaussian-like likelihood function was used to model the floral opening process and a MCMC sampler was used to estimate the peak timepoint. Each RIL had two independently inferred peak floral opening times, from the two blocks of the experiment. The high similarity between replicates verified the robustness of the Bayesian inference protocol. Our method is a hybrid workflow that processes time-stamped high-throughput phenotyping data by feeding raw image data through a machine learning module and a Bayesian inference module in a sequential manner. This modular approach harnesses the respective strengths of the two procedures independently and provides multiple advantages. This flexible workflow can be adapted to different experimental designs and phenotyping goals with only minor changes. Any phenotype that can be scored using time-stamped images could benefit from adopting this workflow with custom likelihood functions based on the biological nature of the phenotype. For example, one could readily substitute with a sigmoid likelihood function for modeling for cumulative phenotypes such as plant height and canopy metrics. Another advantage this workflow has over exclusively-machine-learning-based procedures is that the addition of a specified likelihood function results in interpretable models with biologically meaningful parameters suitable for downstream analyses.

The timepoint at which each RIL reaches peak floral opening varied from early morning to early afternoon in a continuous fashion during the days that the RILs were flowering (Figure 9). QTL mapping identified two loci significantly associated with floral opening time. Each allele from the late flowering parent Armenian999 on the two QTLs contributed additively to a later floral opening phenotype (Figure 11). In contrast to the extensively studied initiation of flowering, floral opening time has been little studied (van Doorn and van Meeteren, 2003; van Doorn and Kamdee, 2014) and not to the locus level (Nitta et al., 2010). This study reports the first genetic loci associated with the regulation of floral opening time.

The largest effect QTL, qDFO2.1, collocates with a QTL that is associated with multiple bolting, budding and flowering time traits in lettuce (Lavelle, 2009). Two genes within qDFO2.1 with non-synonymous variants between the parents have been shown to be involved in the regulation of the circadian clock and initiation of flowering in Arabidopsis (Table 3); these include an ortholog to Arabidopsis UBIQUITIN SPECIFIC PROTEASE 12 (UBP12) and UBP13. *At*UBP13 and *At*UBP12 are ubiquitin-specific proteases capable of rapid, posttranslational regulations of diverse cellular processes in Arabidopsis (Cui et al., 2013). The other candidate gene within qDFO2.1 is orthologous to Arabidopsis CHC1 (alternatively known as BAF60), which is involved in transcriptional activation and repression of flowering regulation genes by chromatin remodeling (Jégu et al., 2014). The absence of floral initiation or circadian clock orthologs within qDFO8.1 suggests that there are separate regulatory mechanisms for floral opening time besides these well studied pathways. The two loci identified in this study provides the foundation for future experiments focused on causal gene identification and functional validation.

The timing of floral opening critically impacts a plant’s survival and reproduction in its community (Kehrberger and Holzschuh, 2019). Our study identified two genetic loci determining natural variation in the regulation of floral opening time in lettuce. This raises the question of the evolutionary pressures for diversity in the trait. Variation in floral opening time may be important in synchronizing floral opening with maximum activity of local pollinators. Thermal constraints on flight activity may limit the pollinating effectiveness of insects; each species of pollinating insect has a microclimatic window within which foraging flight can be sustained (Corbet et al., 1993). The late-blooming parent of the mapping population, Armenian999, is an *L. serriola* accession collected from the cold, wet mountain area of Armenia. In contrast, the early-blooming parent, PI251246, is a landrace accession originating in Egypt. The differential floral opening habits might therefore have evolved in adaptation to different pollinator activities in their respective native environments as has been shown for *Saxifraga oppofitifolia* in alpine elevations (Gugerli, 1998). It would be interesting to investigate whether there is a correlation between floral opening time of diverse lettuce accessions and the climate of their native habitats.

In this study, the combination of machine learning image processing and Bayesian modeling was proved to be highly effective in processing and analyzing time-series aerial images of the field experiment. This versatile framework can be readily adapted to other projects aiming to take advantage of the speed and mobility of drone imaging technologies. Customization in our workflow in choosing suitable machine learning algorithms and Bayesian likelihood functions can enable detection and modeling of phenotypes on the time dimension in other areas such as ecology and population genetics.

## Methods

### Time lapse video and photography

The video was generated with shots at 3 second intervals (3,600 intervals in total), taken with a Canon G15 camera using Canon Hack Development Kit intervalometer script (http://chdk.wikia.com/wiki/CHDK) on June 7^th^, 2014 in the field at Davis, CA. Individual photos were compiled into 30 fps movie clip using PhotoLapse 3 (Version 1.0, S. van der Palen; http://home.hccnet.nl/s.vd.palen/). Close-up photographs of flowers of four asynchronously flowering RILs of the Armenian999 x PI251246 RIL population and the parents were taken at 1-hour intervals on June 7^th^, 2020 using a Canon EOS 50D DSLR Camera. The photographed plants were grown in a screenhouse in Davis, CA.

### Mapping population and field design

Two-hundred and thirty-six F6 RILs had been developed from crossing the *L. serriola* accession Armenian999 with the *L. sativa* landrace PI251246 (M.-J. Truco, unpublished). The 236 RILs, both parental lines, and two controls, *L. sativa* cv. Salinas and *L. serriola* accession US96UC23, were grown in summer 2019 at the Department of Plant Sciences field facility in Davis, CA. The plants were seeded on May 6^th^, 2019 and transplanted into 40-inch-wide raised beds in the field on June 5^th^, 2019. Each raised bed consisted of two rows; every other bed was left empty to allow field access throughout the growing season. The experiment had two complete randomized blocks, each consisting of 240 plots to accommodate the 240 genotypes. Within each block, eight individuals of each RIL or parent were planted into one 10 ft x 1 row plot. The blocks were arranged along the direction of the furrow irrigation system to control for variations attributable to water availability.

### Weather data

Weather data for the dates of the drone flights were collected from the National Centers for Environmental Information website (https://www.ncdc.noaa.gov/) for the University Airport, CA weather station (Station ID WBAN:00174, GPS coordinates 38.533°, −121.783°). The weather station was less than 500 m away from the farthest corner of the experimental field.

### Phenotyping by remote sensing

Seven ground control points were set up in the field, four near the corners and three along the field’s East–West centerline. GPS coordinates, with an accuracy within a few centimeters, were recorded using a handheld data collector (Trimble Geo 7x Series, Trimble Inc., Sunnyvale, CA). These coordinates were used in processing drone images to ensure that images collected at different times and dates aligned perfectly with one another.

A MicaSense RedEdge multi-spectral camera was mounted on a DJI Matrice100 drone. The camera captured images at five wavelengths: blue (475 nm center, 20 nm bandwidth), green (560 nm center, 20 nm bandwidth), red (668 nm center, 10 nm bandwidth), red edge (717 nm center, 10 nm bandwidth), and near-infrared (840 nm center, 40 nm bandwidth). In this study, only the blue, green, and red wavelengths were used for flower identification. The drone was flown over the experimental field at 9 am, 11 am, 1 pm and 3 pm on July 1^st^, 2019, and 10 am, 12 pm, 2 pm and 4 pm on July 9^th^, 2019. The sky was cloudless on both days; daily minimum and maximum temperatures were 13.9°C–31.1°C and 13.9°C–28.9°C; sunrise was at 5:46 am and 5:50am, respectively. A DJI GS Pro app was used to plan and execute the flight. The drone flew at 15 m above ground, and images were taken at a frequency that ensured 85% front- and side-overlaps between each pair of adjacent images. A MicaSense calibration panel was used for automated adjustment of the reflectance spectra. Raw images from the camera were stitched and processed with the Pix4DMapper Pro photogrammetry software to generate orthomosaic maps of surface reflectance at 1 cm spatial resolution. On average, 2,309 raw images were generated per time point, and 2,181 raw images were used to assemble each five-spectrum field map. With the reconstructed maps, the borders of individual plots were manually determined using the software ArcMap.

### Machine learning classification of floral pixels

In order to train a machine learning model that could accurately identify floral pixels from a field image, pixels of flowers, vegetative bodies, and bare ground were randomly sampled and manually labeled from all images based on visual interpretation. A total of 1,569 floral pixels, 1,681 vegetative pixels, and 1,557 ground pixels were labeled (Supplementary Table S1). The HSV values of the sampled pixels were extracted. Half of the pixels (2,404) were randomly selected to be used to train five machine learning models, linear discriminant analysis, k-nearest neighbor, SVM, random forest, and classification and regression tree, using R package “caret” (Kuhn, 2008), for floral pixel identification. A 10-fold within-sample cross-validation test and an out-of-sample validation test with the remaining half of the HSV dataset were performed to compare the prediction accuracy of the machine learning models.

The best performing machine learning model, the SVM model, was trained using HSV values of all 4,807 sampled pixels and used to predict floral pixels for all field images. Once the coordinates of all predicted floral pixels were determined, field images were reconstructed to reflect the floral state of each pixel. A polygonal shapefile delineating the borders of all plots was superimposed on the reconstructed field images to extract the floral pixel counts within each of the 480 plots at each time point. Each plot’s daily maximum floral pixel count was calculated and the plot-level floral pixel counts were normalized by dividing the count number at each time point by the plot daily maximums.

### Bayesian inference of peak floral opening time

A Gaussian-like likelihood function was used to describe the fluctuation of plot-level floral pixel counts of plot *k* (*Y_k,t_, k = 1, 2, …, 480*) at any given time point (t) throughout the course of a day:

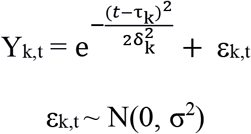

The function peaks at Y = 1 when t = τ_k_. τ_k_ is the center of the symmetric, bell-shaped curve; it describes the time point at which plot *k* reaches its daily global maximum floral opening. Another parameter, δ_k_^2^, determines the spread of the curve, and hence describes the duration of floral opening in plot *k*.

The following weakly regularizing priors were chosen:

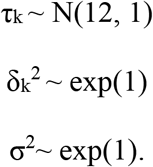

Four 2,000-iteration Markov Chain Monte Carlo (MCMC) were used to sample from the posterior distributions of τ_k_, δ_k_^2^, and σ^2^ using R package “rethinking” (McElreath, 2016). After 2,000 sampling iterations, plots whose MCMC for τ had effective sample sizes below 50 were fed through the modeling pipeline for a second iteration to account for possible poor fitting due to suboptimal initialization. Point estimate of the posterior distribution of τ_k_, 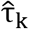, was used as estimated peak FOT for plot *k*. Block effect was assessed using fixed effect simple linear regression. The Euclidian mean between the peak FOTs of the two replicates were used as the phenotypic values for genetic mapping.

### Genotyping and QTL analysis

For DNA extraction, approximately 30 seeds per genotype were placed in a 2 mL Eppendorf Safe Lock tube along with one stainless steel bead (Qiagen Cat. No. 69989), frozen in liquid nitrogen, and ground to a powder in a Qiagen TissueLyser. Seven hundred microliters of 5 M guanidine thiocyanate in 20 mM Tris-HCL (pH 6.75) was added to tissue powder, vortexed until homogenized and spun in microcentrifuge (RT) for 5 min at 14,000–20,000 g. After centrifugation, 600 μL of clear lysate was transferred to DNA binding plates (Epoch Life Sciences EconoSpin™ 96 well) stacked over a 1 mL collection plate and centrifuged 5 minutes (RT) at 1,300 g. Flow through was discarded. DNA binding plate was incubated with 600 μL of liquid for 4 minutes at RT and centrifuged at 1,300 g for 5 minutes sequentially with PB buffer (Qiagen Cat. No. 19066), followed by PE buffer (Qiagen Cat. No. 19065), and then with two 80% EtOH washes. DNA plate was dried in centrifuge over paper towels for 5 minutes at 2,000 g. DNA was eluted from binding plate into new collection plate after 5-minute incubation in 60 μL of 10 mM Tris-HCL (pH 8.0) at 65°C. DNA was quantified using Qubit. DNA from parental lines and segregating individuals was digested using *AvaII* to reduce the genome complexity of the samples (Sandoya et al., 2019). Individual samples were barcoded, pooled, and genotyped by sequencing using paired-end 100 bp Illumina HiSeq4000. The parental lines, Armenian999 and PI251246, were also whole-genome-shotgun sequenced using paired-end 150 bp and 100 bp Illumina HiSeq4000 to 29x and 17x coverages, respectively. Sequencing results were de-multiplexed using GBSX software in the demultiplex mode (Herten et al., 2015). All reads were mapped to the *L. sativa* reference genome v8.0 (Reyes-Chin-Wo et al., 2017) using bwa-mem (Li, 2013) and variants were called using FreeBayes (Garrison and Marth, 2012). SNPs called against the reference that were polymorphic between the two parental lines with a quality score greater than 20 and with fewer than 20% missing data across all RILs were used to construct a genetic map using the software LepMap3 with 20 cM as the cutoff threshold for linkage groups and the significance level at p-value = 10^-6^. One representative SNP from each genetic bin was selected for QTL mapping. Linkage group numbers were determined by the chromosomal location of the markers relative to the reference genome. Heritability of the phenotype was estimated using mixed effect modeling with R packages “synbreed” (Wimmer et al., 2012) and “sommer” (Covarrubias-Pazaran, 2016) using block-level phenotype data. QTL analysis was performed using 2,677 SNP markers, each representing distinct genetic bins. The R package “qtl” was used for interval mapping, 10,000-iteration permutation test, and QTL effect analysis (Broman et al., 2003).

### Candidate gene identification

Single nucleotide variants, insertions, deletions, stop-loss variants and stop-gain variants were identified between the parental lines using software ANNOVAR (Wang et al., 2010). Genes were filtered for non-synonymous variations in coding sequence between the parental lines. Orthofinder (Emms and Kelly, 2015) was used for genome-wide prediction of lettuce orthologs of *Arabidopsis thaliana* genes.

## Acknowledgements

We thank J. Emerson for greenhouse and field assistance, A. Vargas for DNA and GBS library construction, D. Feinberg for assistance with raw image processing, and H. Xu for data submission to NCBI Sequence Read Archive (SRA) database. This research was funded by an NSF Graduate Research Fellowship to RH and a USDA NIFA Specialty Crop Research Initiative (SCRI) Grant # 2015-51181-24283 to RWM.

## Data Availability

GBS data of the RILs and WGS data of the parents are available on the NCBI SRA database under BioProjects PRJNA642889, PRJNA510128, and PRJNA478460, respectively. Scripts used in the study for machine learning and Bayesian inference are available on GitHub at https://www.github.com/rkbhan/FloralOpening. GPS-anchored aerial image data are available on HydroShare at https://www.hydroshare.org/resource/1c5855dbeb3c49a8b5779300550e08f1/.

